# The regulatory function of dIno80 correlates with its DNA binding activity

**DOI:** 10.1101/519959

**Authors:** S Jain, J Maini, A Narang, S Maiti, V Brahmachari

**Affiliations:** University of Delhi; Institute of Genomics and Integrative Biology

**Keywords:** Chromatin remodelling, *Drosophila*, Ino80, DNA Binding, Consensus sequence, Non-canonical complex, Transcription regulation

## Abstract

The INO80 complex, including the Ino80 protein, forms a highly conserved canonical complex that remodels chromatin in the context of multiple cellular functions. The *Drosophila* homologue, dIno80, is involved in homeotic gene regulation during development as a canonical Pho-dIno80 complex. Previously, we found that dIno80 regulates homeotic genes by interacting with epigenetic regulators, such as polycomb and trithorax, suggesting the occurrence of non-canonical Ino80 complexes. Here using spectroscopic methods and gel retardation assays, we identified a set of consensus DNA sequences that DNA binding domain of dIno80 (DBINO) interacts with having differential affinity and high specificity. Testing these sequences in reporter assays, showed that this interaction can positively regulate transcription. These results suggest that, dIno80 has a sequence preference for interaction with DNA leading to transcriptional changes.

**SIGNIFICANCE:** The chromatin remodeling proteins control gene expression by nucleosome sliding and exchange. They are known to function as multi-subunit complexes recruited to chromatin by transcription factors or histone modification readers. Here, we report a sequence specific binding potential for the chromatin remodeler, dIno80. We have carried out *in vitro* studies with DNA binding domain of dIno80 to elucidate its sequence specific DNA binding. We have also showed that this binding can regulated reporter gene expression in *Drosophila* cells. Our results suggest a non-canonical role of Ino80 in transcriptional regulation.

## INTRODUCTION

In eukaryotes, the chromatin provides the platform for the interaction of various regulatory factors to control transcription temporally as well as spatially. This regulation operates through post-replication modification of DNA (1–2) and the post-translational modification of histones (3–4). The most significant epigenetic marks are the covalent modifications of DNA and histone proteins associated with DNA, in particular DNA methylation (1) and histone tail modifications, such as acetylation, methylation, phosphorylation, ubiquitination, ADP ribosylation, and glycosylation (5). These marks are recognized by the ATP-dependent chromatin remodeling complexes which are multi-protein complexes that modulate chromatin dynamics (6).

All known ATP-dependent chromatin remodeling enzymes belong to the helicase superfamily 2 (SF2), as their ATPase domain harbours seven motifs that are characteristic of helicases (7). Using a phylogenetic approach these proteins are divided into subfamilies based on the architecture of their ATPase domain and the presence of motifs outside the ATPase domain. The well-known subfamilies are: SWI2/SNF2 (switch 2/ sucrose-non-fermenting 2)-related proteins, CHD (chromodomain-helicase-DNA binding) family, INO80/SWR1 family and ISWI (imitation switch)-related protein family (8–10). With a few exceptions, these ATPases are genetically non-redundant, with mutations in these genes often having early embryonic and/ maternal-effect phenotypes. ATP-dependent chromatin remodeling proteins are functionally diverse and are involved in a variety of cellular functions like telomere maintenance, chromosome segregation, checkpoint control, DNA replication, DNA damage response and transcriptional regulation by virtue of multiple interactions (11). These enzymes function in a DNA dependent manner and are recruited to multiple sites on the genome through the interaction with DNA binding proteins. One such DNA binding protein is the transcription factor Pleiohomeotic (Pho) in *Drosophila melanogaster* (12).

The INO80 protein was identified in a screen for mutants for inositol biosynthesis (13). As evidenced by its role in phospholipid biosynthesis, yINO80 (yeast INO80) brings about the positive as well as negative regulation of several promoters (14–15). The INO80 ATPase is a member of the SNF2 family of ATPases and functions as an integral component of a multi-subunit ATP-dependent chromatin remodeling complex. The INO80 subfamily is conserved from yeast to higher organisms and there the well-known INO80 complex has highly conserved subunit composition (16–18). In previous work, the human and the *Drosophila* homologues of Ino80 have been identified (16–17). Klymenko *et al*., (19) identified dIno80 as a part of a complex that contains Pho (Pleiohomeotic) in *Drosophila*. The INO80 homologues from *Drosophila* as well as human was shown to interact with the factors that are similar to those known for the well characterized INO80.Com in *Saccharomyces cerevisiae* (18). This can be considered as the canonical complex. Some of the proteins present in the complex that have attracted attention are the Actin Related Proteins, ARPs. Among the ten ARPs known in *S. cerevisiae*, six (Arp 4-9) are known to be associated with chromatin regulating complexes (20, 21). However, both the *Drosophila* and the human canonical complex of INO80 contain fly specific and mammalian specific proteins respectively, in addition to the conserved members like SNF2 family ATPase, AAA^+^ ATPases Tip49a and Tip49b, ARPs and Ies2 and Ies6 proteins (22).

The role of dIno80 in *Drosophila* and its interaction with PcG-trxG complex was demonstrated in previous reports from our lab (23–24). The effect of INO80 complex on transcription is linked to the function of the transcription factors yin yang 1 (YY1) in mammals and Pleiohomeotic (Pho) in *D. melanogaster*, which are part of the INO80 complex (12, 19, 25). In addition to Pho/YY1, the INO80 complex contains two AAA+ ATPases (ATPases associated with variety of cellular activities) referred to as RUVBL1 (in mammals and Reptin in *D. melanogaster*) and RUVBL2 (in mammals and Pontin in *D. melanogaster*). Ino80, along with Reptin and Pontin, has also been reported to be involved in immunoglobulin class-switch recombination by regulating the cohesin activity (26). The proteins Reptin and Pontin have well documented role in embryonic development as members of PcG-trxG protein complexes (19, 28). The Pho independent function of dIno80 has been demonstrated (24). In addition, it is known that loss of function mutants of *dIno80* show embryonic lethality, implicating the essential role of dIno80 in developmental gene regulation (23). The INO80 subfamily functions in diverse cellular processes and is reported to interact with promoters of pluripotency genes, dependent on Oct4/WDR5 binding (22, 28). In an independent investigation, we have demonstrated that the hIno80 exerts transcriptional regulation through its DNA binding domain (DBINO) (17, 29). This study identified a consensus sequence motif (GTCAGCC) through which the hIno80 protein interacts with chromatin.

In the light of the interaction of dIno80 with polytene chromosomes (24) and the known DNA binding activity of human INO80 (29), we examined the DNA binding properties of dIno80 with the consensus sequence identified earlier for hINO80 and its effect on gene expression. Since, we find that dIno80 interaction leads to activation of target gene transcription, we carried out a detailed analysis of the binding properties of dIno80. We identified a domain, approximately 250 amino acids in length (DBINO domain), that shows DNA binding activity *in vitro.* The kinetic analysis indicates that dIno80 distinguishes between different binding sites on the genome. We suggest the dual function of dIno80 in transcription regulation in *Drosophila,* both as a DNA binding developmental regulator and a DNA-dependent ATPase.

## MATERIALS AND METHODS

### Purification of recombinant DBINO domain of dIno80

The 5238 bp cDNA clone (Flybase Id: FBcl0292012) of dIno80 was obtained from Drosophila Genomics Resource Center, Bloomington. The cDNA from embryonic mRNA was also synthesized. The 800 bp region (838 to 1605bp), including the DBINO (364-489 amino acid) domain along with the flanking region in dIno80 cDNA was PCR amplified using primers dDBINO-256-MAL-Fp: 5’-ATAGGATCCTCAGAATCCGGCG-3’ and dDBINO-256-MAL-Rp: 5’-CCCAAGCTTTCTTTGCATAGCTGC-3’. The region corresponds to 279-535 amino acid in the dIno80 coding sequence. The forward primer was designed to contain a BamHI site and the reverse primer contained a HindIII site. The resultant 800 bp amplicon was digested with BamHI and HindIII and directionally sub-cloned into pMAL-c2x expression vector (New England Biolabs, Massachusetts, United States). The protein was expressed in BL21 (DE3) strain of *Escherichia coli* at 16°C for 12 hours in LB medium supplemented with 0.2% glucose following induction with 0.5mM IPTG. The recombinant DBINO domain containing protein (~80kDa) was purified using the amylose affinity column (New England Biolabs, Massachusetts, United States) and detected using the anti-MBP-HRP antibody (NEB). The bound protein (~42kDa, 360 amino acids) was released from the MBP tag by cleavage with Factor Xa (New England Biolabs, Massachusetts, United States) for 24 hours at 4°C in 20mMTris pH 7.2, 100mMNaCl and 100mM CaCl_2_. The protein at this stage was concentrated and exchanged into 20 mM Tris pH 7.5 and 50mMNaCl. This sample was analysed by mass spectroscopy analysis in Dr. Suman Thakur’s Lab, Center for Cellular and Molecular Biology, Hyderabad.

### FID (Fluorescence Intercalator Displacement) assay

FID assay is based on the displacement of ethidium bromide from the double-stranded DNA oligo (30). The steady-state fluorescence emission intensity measurements were performed with a Spex Fluoromax-4 spectrofluorimeter. Excitation and emission slits with a nominal band-pass of 5 nm were used for all measurements. Background intensities of samples without protein were subtracted from each spectrum to avoid any contribution from solvent, Raman peak, and other scattering artefacts. The fluorescence titration was performed using fixed concentration of dsDNA (1 μM) and varying concentrations of DBINO protein (0.2–5 μM). The DNA: EtBr ratio was kept as 1:0.5. The dsDNA oligo was prepared by annealing the single stranded oligo and its reverse complement oligo at 100 μM concentration in the annealing buffer (10mMTris-Cl pH-7.5-8.0, 50mMNaCl and 1mM EDTA). The oligo sequences are given in Table S1. All buffers used were filtered through 0.2 micron filters. The mixtures were stirred continuously in a 1.0-cm path length cuvette to ensure homogeneity and all titrations were carried out at 20°C. The EtBr-DNA sample was excited at 480 nm in 20mM HEPES pH7.0 and 50mMNaCl buffer. The emission was measured at 500–700 nm wavelength range. Further an aliquot of protein was added until there was no further change in fluorescence intensity in emission spectrum. The change in fluorescence intensity at 600 nm (λ_max_ for EtBr) was then used to interpret the data. The data was analyzed using the double reciprocal plot.

### Electrophoretic Mobility Shift Assay

The oligonucleotide sequence having three Ino80 binding motif was cyanine (Cy3) labeled at the 5’ end. The reverse complement of the same sequence was also synthesized. The list of oligonucleotide sequences is given in the Table S1. The oligos were annealed in the annealing buffer (10mMTris-Cl pH-7.5-8.0, 50mMNaCl and 1mM EDTA). The nuclear extract was prepared from larval tissue (~20 larvae) of *Canton-S* stock using the CelLytic™ NuCLEAR™ Extraction Kit (Sigma, Missouri, Unites States). The nuclear extract was taken in different amounts and incubated in the presence of 1X binding buffer (250mM HEPES, 10mMKCl, 1mM EDTA, 10mM DTT, 1mM PMSF and 100mMNaCl), 1 μg Poly (dI-dC), 1 μg tRNA and 10% glycerol for 10 minutes at room temperature. The Cy3 labeled (2-10 picomoles) double-stranded oligonucleotide was added to the reaction mixture and kept for 20 minutes at room temperature. The mixture was analyzed on 5% acrylamide-bisacrylamide (29:1) gel in 0.5X Tris-borate-EDTA (TBE) buffer pH8.0 with 4% glycerol. Following electrophoresis at constant voltage (80V), the gel was scanned on Typhoon phosphoimager (Typhoon FLA TRIO, General Electric Healthcare, Little Chalfont, United Kingdom). The probe to protein ratio was considered in terms of molar ratios, to have an idea of the number of molecules of the two interacting components. For this empirically we assumed an average molecular weight of the protein as 110 kDa. For the competition experiments, specific amount of unlabeled (cold) oligos in the ratio of 1:1 to 1:10 was added in different reaction sets for analyzing the specificity of interaction. Similarly, the cold non-specific probes were also analyzed. For the variants of Hs.con, the oligos listed in Table S1 were used.

### Cell Culture and Reporter Assays

The S2 (Schneider 2) cell lines have been derived from a primary culture of late stage (20-24 -hours) *Drosophila melanogaster* embryos (31). S2 cells were cultured in Schneider’s *Drosophila* medium (Gibco^®^, Life technologies^™^, California, United States) containing 10% FBS (Fetal Bovine Serum, qualified, US origin, Gibco^®^, Life technologies^™^, California, United States) and Penicillin-Streptomycin (10,000 U/mL- Gibco^®^, Life technologies^™^, California, United States) at a final concentration of 50 units penicillin G and 50 μg streptomycin sulphate per milliliter of medium in 25°C incubator (Memmert^™^ GmbH, Schwabach, Germany). For nuclear extract, the S2 cells were transfected with 450 picomoles of siRNA for dIno80 and control siRNA (SIC001, Sigma, Missouri, Unites States) at 50% confluency in a T-75 flask. For carrying out the knockdown experiment, we used two different pools of duplex siRNAs for dIno80 (dIno80 (NM_169854)-duplex1: 5’-CGAGGAAAUUGAAAUGAAA-3’, dIno80 (NM_169854)-duplex2: 5’-GACUCCAUAUGAUCUUGAA-3’), which were subsequently mixed. The cells were maintained in incomplete media in 25°C incubator for 8 hours after transfection and were then maintained in complete media for 72 hours hours. The cells were, then, harvested to prepare nuclear extract using the CelLytic™ NuCLEAR™ Extraction Kit (Sigma, Missouri, Unites States). For luciferase reporter assays, 2.2×10^5^ cells were transfected with 1,000 ng pGL3-promoter luciferase vector or cloned constructs (28), 10ng of hRLuc pGL4.75 control vector in 24-well plate format using FuGENE® HD transfection reagent (Promega Corporation, Wisconsin, United States). The cells were maintained in incomplete media in 25°C incubator for 8 hours after transfection and were then maintained in complete media for 48 hours. The cells transfected in 24-well plates were assayed post-transfection for relative luciferase activity using Dual-Luciferase Reporter (DLR™) Assay System (Promega Corporation, Wisconsin, United States) according to manufacturer’s protocol. The relative luciferase activity was calculated by normalization of Firefly luciferase counts to Renilla and compared with the control sample. We performed transfection experiments using 24 picomoles of siRNA pool. The control siRNA was procured from Santa Cruz Biotechnology, Texas, United States (sc-37007) and Sigma, Missouri, Unites States (SIC001). The RNA was isolated using TRIzol (Invitrogen, California, United States) method 48 h post transfection. The cDNA was synthesized utilizing the First Strand cDNA Synthesis Kit from Thermo Scientific, Massachusetts, United States (Cat # K1612), using oligo-dT primers. The qPCR experiments were carried out in an Applied Biosystems machine (ABI 7300), using FastStart Universal SYBR Green Master (Rox) Roche Diagnostics GmbH, (Mannheim, Germnay) with cDNA as the template. The cycle threshold (C_t_) values obtained for test gene from siRNA treated cells (C_t_ siRNA, test) and untreated cells (C_t_ wild type, control) were normalized to the endogenous control, *Rpl32* from the respective cell types. The normalized values were compared. We analysed the relative gene expression, using the 2^−^ΔΔCT method (32). The primers used are as follows: *ino80* Fp: 5’-GGGAACGTCTTCACCCCTAC-3’, *ino80* Rp: 5’-CAGGACACTCCAGGCGATTG-3’, *rpl32* Fp: 5’-AGCATACAGGCCCAAGATCGTGAA-3’, *rpl32* Rp: 5’-TCTGTTGTCGATACCCTTGGGCTT-3’. This was performed in three biological replicates each in triplicate. The test of significance used is Student’s t-test (two-tailed).

### *In silico* analysis and gene ontology classification

The binding peaks for Ino80 were retrieved from the public database. The dIno80 ChIP-chip data is available on gene expression omnibus (Accession number GSE32404) (33) The Pho ChIP-seq data was retrieved from Schuettengruber *et al*., (34). The overlapping and unique regions with pho and dIno80 binding sites were extracted using bedtools. Further the genes mapping within ±2.0kb of the peaks were identified using UCSC table browser (*Drosophila* reference genome, release - dm6, August 2014). These genes were grouped into overlapping and unique Pho and dIno80 putative targets. The Gene Ontology classification at the level of biological process of the genes was performed using PANTHER v12 tool (35).

### *Drosophila* stocks

Wild-type (*Canton-S*) *D. melanogaster* was used in all the experiments and were maintained on corn agar medium at 25 °C.

## RESULTS

### DNA binding activity of dIno80

The human and the *Drosophila* homologues of INO80/Ino80 proteins showed 90% homology, while the DBINO domains have 97% homology. Mendiratta *et al*., (29) reported the interaction of DBINO domain with a specific DNA binding motif 5’ [CA][CA][CA][CG] GTCA[GC]CC 3’. We refer to this motif as human consensus, Hs.con, subsequently in this manuscript. Since the DBINO domain is conserved across the phyla, we examined the DNA binding ability of *Drosophila* homologue, dIno80 (16–17). We cloned and purified the DBINO domain of dIno80 in pMAL-c2x vector yielding a 42kDa protein product (Fig. 1, A and B). The identity of the protein was confirmed using the mass spectroscopy analysis by the detection of unique peptides with at least 5 peaks as shown in the representative spectra for one of the peptides KLGQGSEEDQLR (Fig. 1 C). We identified a total of 11 unique peptides mapping to DBINO domain of dIno80 protein out of which 4 peptides showed 5 peaks.

**FIGURE 1.**
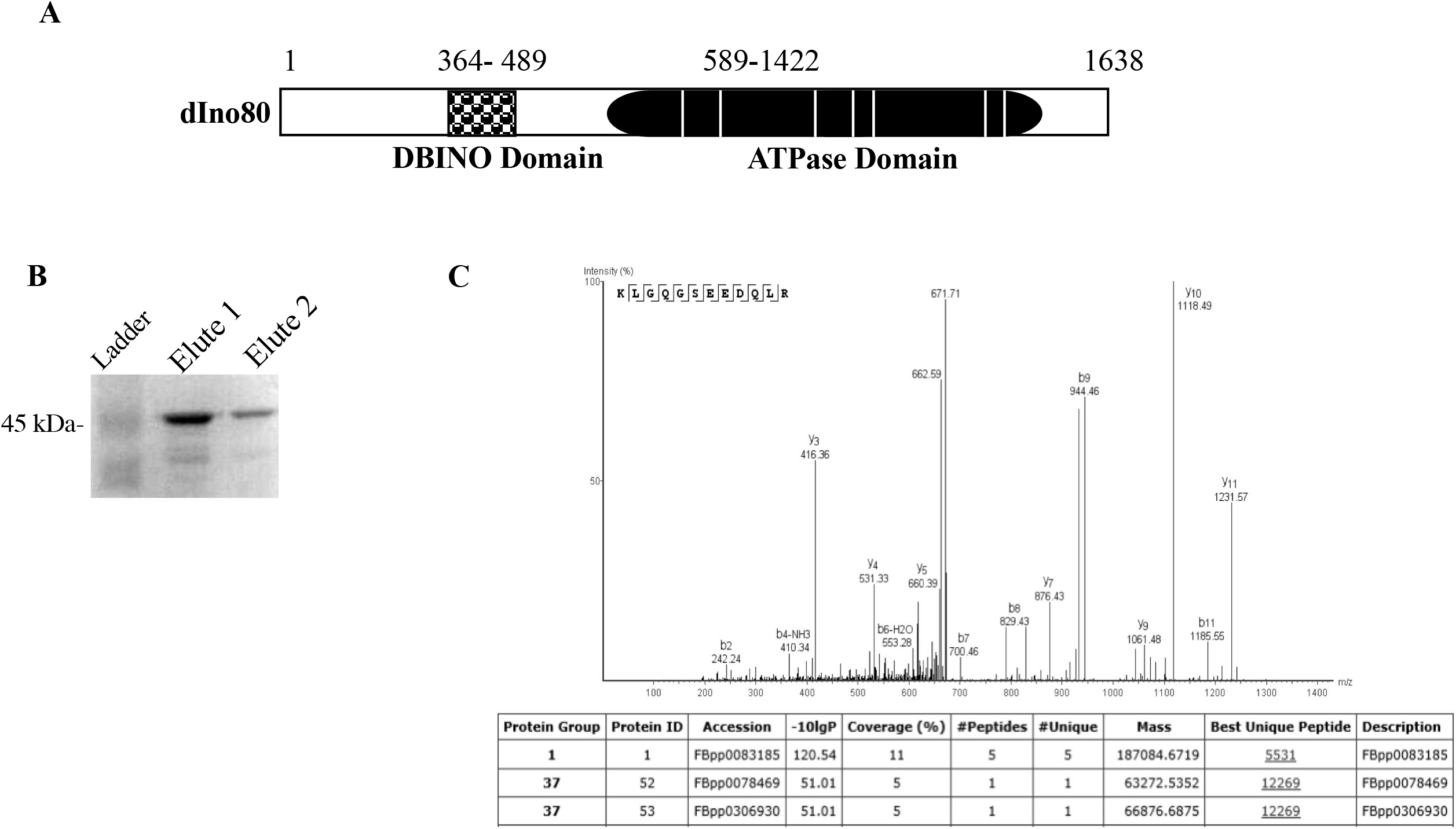
The expression of DBINO domain of dIno80 from the clone. (A) Line diagram showing the relative positions of the putative DNA binding domain (DBINO) in the full length protein. (B) Expression of the cloned and tag-cleaved DBINO domain was visualized by silver staining C) The analysis of the DBINO domain by mass-spectroscopy, the unique peptide (KLGQGSEEDQLR) mapping to the DBINO domain is shown.

The interaction of the purified DBINO domain of dIno80 with Hs.con sequence, was analyzed by Fluorescence Intercalator Displacement (FID) assay (Fig. 2). In FID method, change in fluorescence intensity of EtBr bound DNA is measured as the displacement of DNA-bound ethidium bromide (EtBr) by the protein. A 33-mer oligonucleotide containing 3 repeats of Ino80 binding motif (Hs.con) was used as the specific probe in these reactions. The displacement of the bound ethidium bromide by the DBINO domain showed incremental loss in fluorescence and a well-defined titration curve with the specific oligo, while with the non-specific oligonucleotide, DBINO domain failed to displace the ethidium bromide from the DNA-EtBr complex (Fig. 2, A and B). The titration curve was plotted as the change in the relative fluorescence intensity (I/I_0_) as a function of increasing protein concentration (Fig. 2 B). To calculate binding affinity, the same data was re-plotted as double reciprocal plot with reciprocal of fractional saturation as a function of reciprocal of protein concentration (molar units) and the affinity, K_d_ = 10 μM, was calculated as described by Ham *et al.*, (30) (Fig. 2 C). The low affinity could be because the purified 279 amino acid peptide corresponding to the DBINO domain was used in these assays; while the INO80 complex is the functional form within the cell. Considering this we have carried out the binding assays with nuclear extract as described in the next section.

**FIGURE 2.**
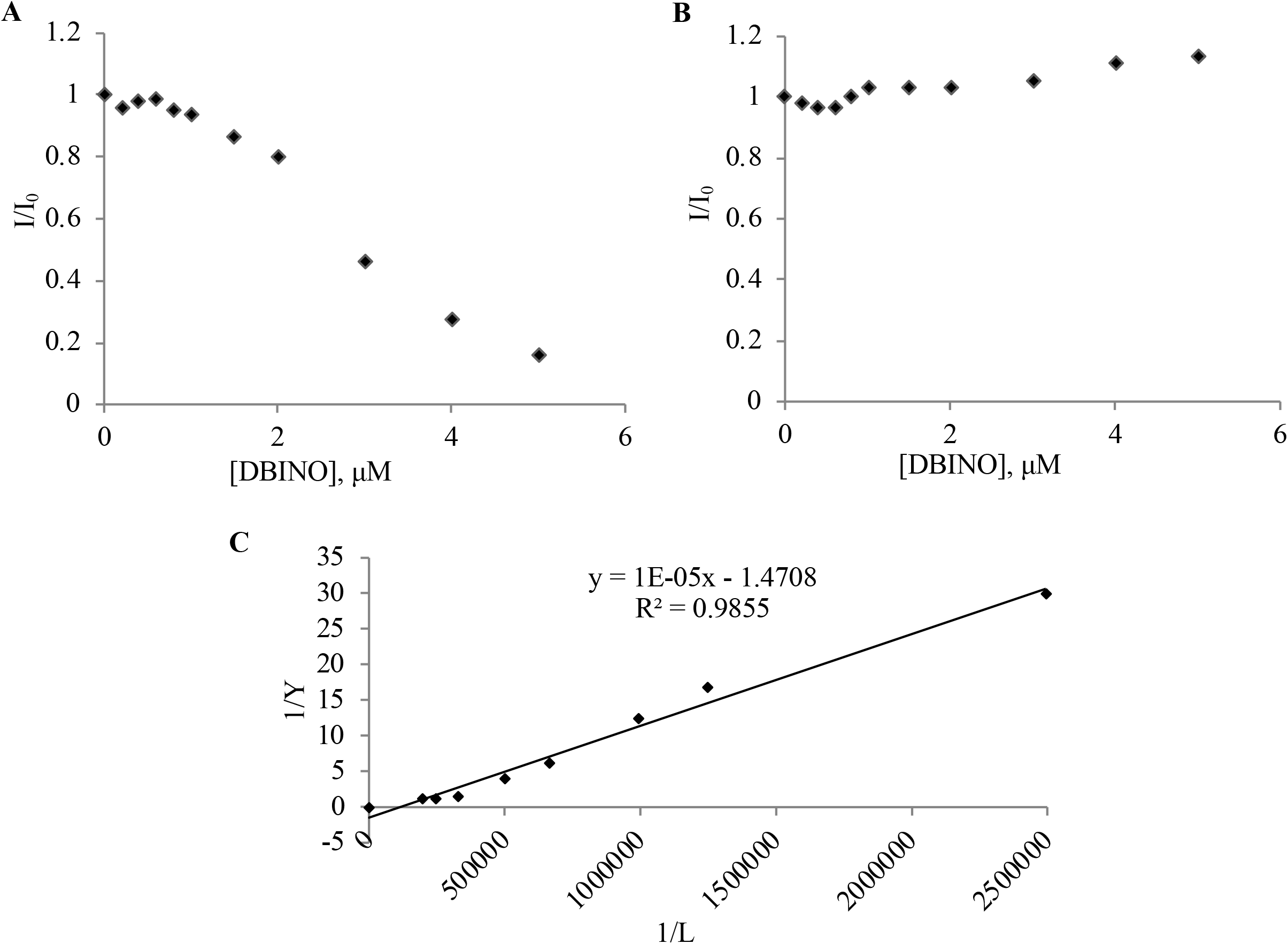
The interaction of DBINO domain of dIno80 with oligonucleotides. The binding was monitored as the displacement of ethidium bromide from DNA by DBINO using fluorescence spectroscopy. The specificity of the interaction is shown by using the specific oligo, Hs.con sequence (A) and the non-specific oligo (B). Ethidium bromide is displaced by the protein only in case of the specific oligo and not the non-specific oligo. The sequence of the two oligos is given in the Supplemental Table S1. (C) The double reciprocal plot for interaction of recombinant DBINO domain of dIno80 with Hs.con-EtBr complex. The reciprocal of change in fluorescence intensity (1/Y) is a plotted against reciprocal of molar equivalent of protein (1/L). The change in fluorescence intensity was calculated as ΔF/ΔF_max_. The slope of the graph is inversely proportional to the K_d_ for binding (Ham *et al*., 2003).

### Analysis of DNA binding activity of the native dIno80 in nuclear extract

We examined the interaction of dIno80 with the predicted Hs.con motif by Electrophoretic Mobility Shift Assay (EMSA) using nuclear extracts from *Canton-S* (Fig. 3 A). The nuclear extract from third instar larvae was used in these assays with Cy3 labeled double-stranded oligonucleotide having three copies of the Hs.con sequence at 1:1 to 1:50 molar ratios of oligo: protein. The amount of oligo used was 2 picomoles which is 0.086μg of DNA and the protein was varied from 0.2-10μg. The binding is detected at a molar ratio of 1:5 and it increases in a concentration dependent manner (Fig. 3 A). The intensity of the bound oligo as well as the extent of shift increases with an increase in protein concentration, as shown by higher shifted bands. Using the densitometric analysis of the gel, the K_d_ for binding was found to be 3.1μM (Fig. 3 B). The specificity of the interaction of the nuclear extract with the DNA probe was tested using boiled nuclear extract, bovine serum albumin as non-specific protein (Fig. S1 A), and proteins remaining in the nuclear pellet as control extracts (Fig. S1 B). None of these samples were able to interact with the Hs.con sequence and hence, no shift was visible on the gel (Fig. S1). Due to the lack of anti-dIno80 antibody we could not carry out supershift assays. In a study involving the ARP containing sub-complex of the Ino80 complex, the authors consider affinities less than 5000nM (5μM) as specific interactions (36).

**FIGURE 3.**
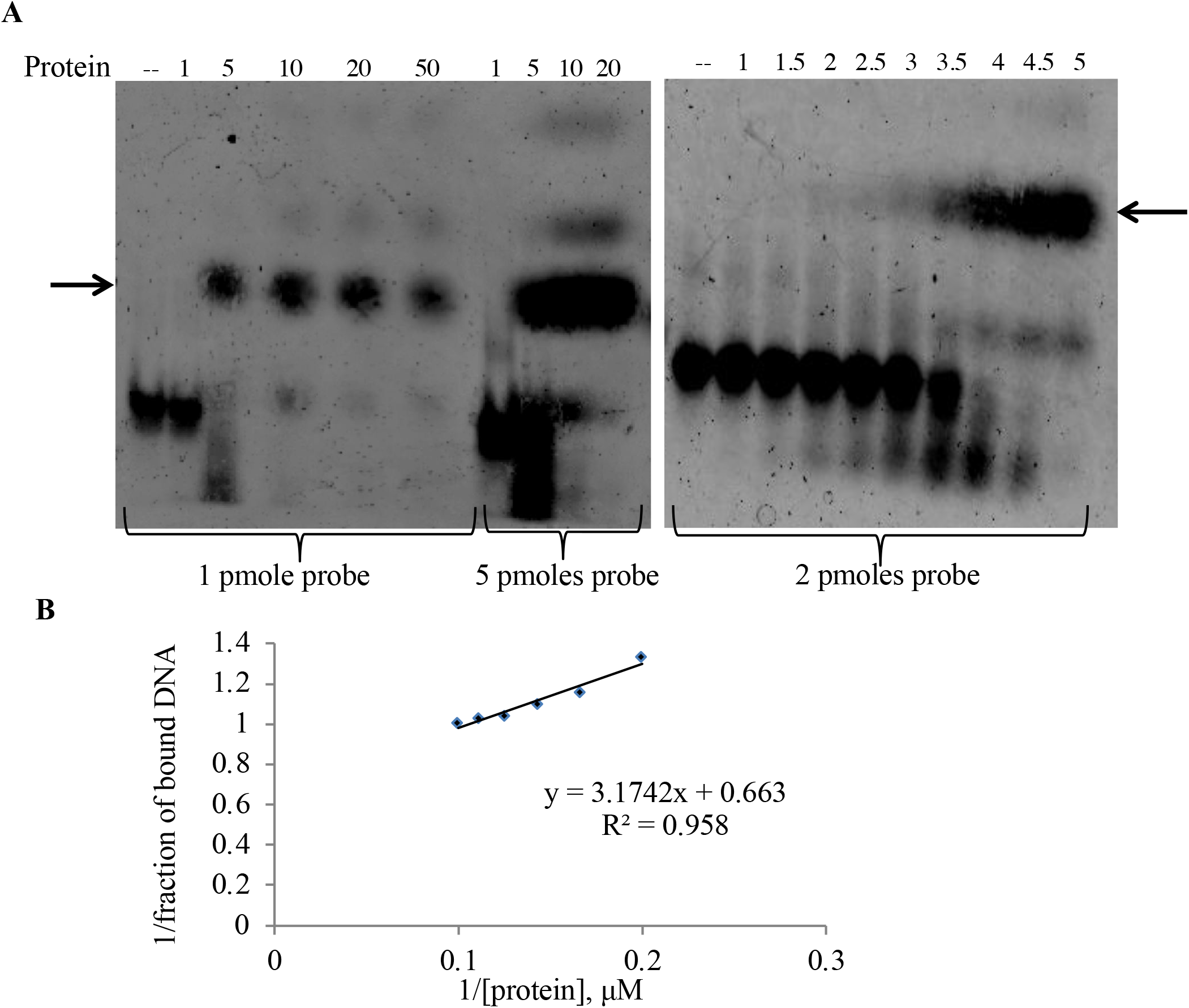
Interaction of Hs.con with nuclear extract from wild type larvae. (A) Electrophoretic mobility shift assay was carried out with Hs.con oligonucleotide end labelled with Cy3. The concentration of the probe was 1,5 and 2 picomoles as grouped. In each group the probe concentration was kept constant and the concentration of the protein (molar ratio) was varied as shown on the top of the lanes, (B) The double reciprocal plot for K_d_ calculation. K_d_ is estimated to be 3.1μM.

The specificity of interaction was also confirmed by comparing the EMSA with the nuclear extract from larvae with heterozygous deletion for *dIno80* and the wild type (*dIno80*^+/−^vs. *dIno80*^+/+^) with increasing probe: protein ratio (1:1-1:100; Fig. 4 A). The copy number of *dIno80* wild type gene is different in the two cases, with deletion mutant having only one functional copy of the gene. We could not use nuclear extract from dIno80 null mutants as they are non-viable. The relative intensity of the retarded band was compared between the wild type and the mutant at the same oligo: protein ratio. The intensity of the band in dIno80^+/+^ was compared with that in *dIno80*^−/+^ at six different ratios of oligo to protein, ranging from 1:1 to 1:100. The extracts from heterozygous mutant larvae showed decreased intensity of the retarded band as seen from densitometric profile (Fig. 4 B). At 1:1 (probe: protein) ratio, no retardation was seen in both genotypes. The retarded band obtained using heterozygous larvae (*dIno80^−/+^*) showed nearly 50% less intensity as compared to wild type larvae (*dIno80*^+/+^) at 1:5 probe:protein ratio. However, at higher molar ratio (1:100 probe:protein), the relative band intensity is almost the same for both the wild type and mutant larval extracts, due to saturation of the probe and the protein complex. The extract is prepared from the heterozygous mutant larvae, having reduction in protein levels by half only. This explains the possible saturation of all the binding sites at the higher molar ratio of probe: protein.

**FIGURE 4.**
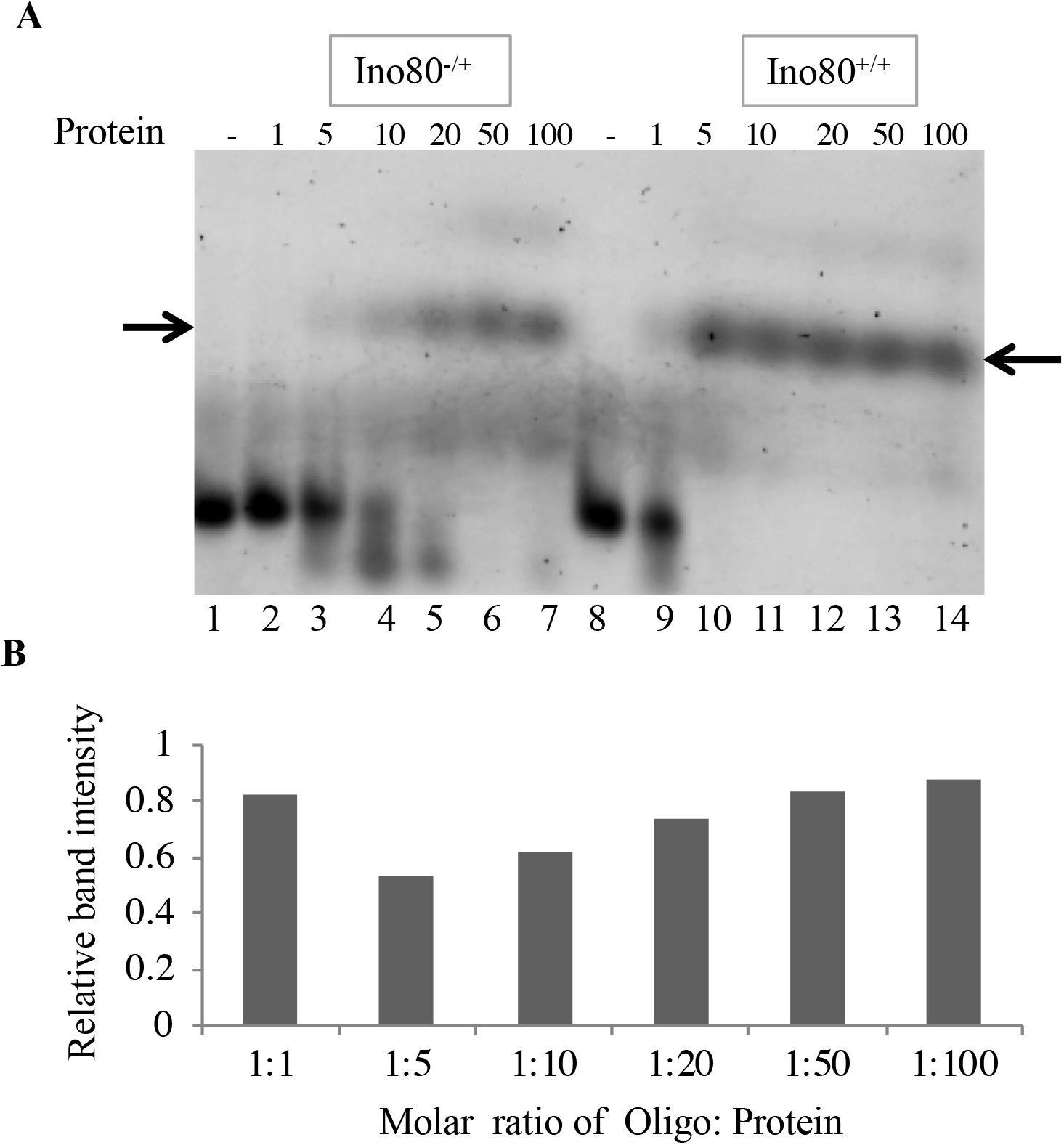
Specificity of interaction of Hs.con with dIno80. (A) Interaction of Hs.con with the nuclear extracts from mutant Ino80^−/+^ larvae (lane 2-7) compared to nuclear extract from wild type (Ino80^+/+^; lane 8-14) at increasing concentration of the protein (X times probe) and a fixed concentration of the probe. (B) Densitometric scan of gel shown in (A); The relative intensity on Y axis is the ratio of the retarded band between Ino80^+/+^ and Ino80^−/+^ at given ratio of probe:protein as indicated on the X-axis.

To determine the specificity of the interaction, we tested the displacement of Cy3-specific oligo by unlabeled (cold) specific and non-specific oligonucleotides in competitive EMSA (Fig. 5 A). The binding of the probe was competed out by the unlabeled specific oligo at a molar ratio 1:5 (Cy3-labeled to unlabeled specific oligo). The displacement was almost 60% at the molar ratio of 1:10 (Fig. 5 B). The non-specific unlabeled oligonucleotide failed to compete out the specific labeled oligo, thus, establishing the specificity of the interaction (Fig. 5 A). The specificity of this interaction was further strengthened by lack of competition by random oligonucleotide containing a scrambled sequence of human consensus motif (11mer) in three biological replicates, which failed to compete out the labeled Hs.con (Fig. 5 C).

**FIGURE 5.**
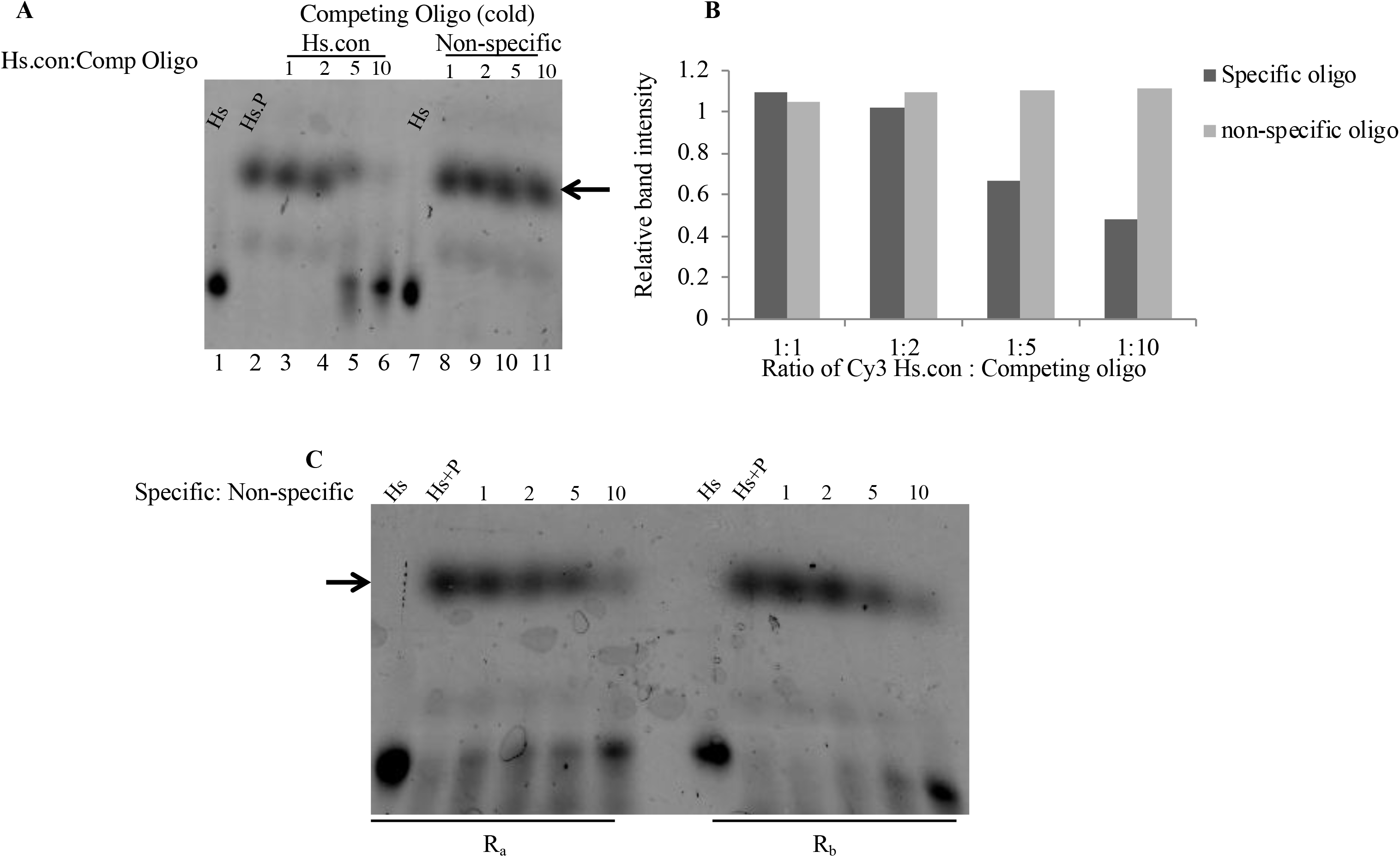
Specificity of interaction of Hs.con with dIno80 using competitive EMSA. (A) Results of competitive EMSA; Cy3 labeled specific oligo and unlabeled competing specific-(lane 3-6) and non-specific oligos (lane 8-11). The competing oligo was varied at different molar ratio relative to the specific oligo as indicated at the top of the lanes in terms of X times the labeled probe. The probe (Cy3-Hs.con) to protein ratio was the same (1:5) in all the experiments. (B) Densitometric scan of gel shown in (A). For this, the ratio was calculated between the retarded band in presence of competing oligo with respect to that of in its absence. In (B), the Y-axis, the ratio of intensity of the retarded band in absence of competing oligo relative to that in its presence has been plotted against the varying ratio of probe: competitor on X-axis. (C) Results of EMSA with Cy3 labeled specific oligo at molar ratio of 1:5 (probe: protein) in all the experiments. The unlabeled competing non-specific oligos (R_a_ and R_b_) were varied at different molar ratios as indicated at the top of the lanes in terms of X times the labeled probe. The 33mer oligos were synthesized **with randomized** sequence repeated thrice. (R_a_ = CCCCCAGTCG; R_b_= CCCCCGTGCTA). The molar ratio used is indicated on the lanes.

To further confirm the specificity of interaction of dIno80 with Hs.con, the nuclear extract from S2 cells transfected with and without siRNA against dIno80 was used. The siRNA knockdown reduced the expression of *dIno80* by 80% (p=0.00007, Student’s T-test; Fig. S2). The normalization control used for qPCR was *rpl32*. The nuclear extract from S2 cells transfected with siRNA against dIno80 failed to show any shift with Hs.con oligo as opposed to the nuclear extracts from wildtype S2 cells or cells transfected with control siRNA duplexes, indicating the specificity of binding of dIno80 (data not shown). These results establish the specificity of interaction between the human Hs.con and the DNA-binding domain of dIno80 (DBINO). The EMSA was performed at five different ratios of oligo: protein ranging from 1:1 to 1:80.

### Regulatory role of the interaction of dIno80 with DNA

To understand the consequence of the binding of dIno80 to DNA on transcriptional regulation, the dIno80 binding motif was cloned in pGL3 promoter vector with firefly luciferase as the reporter (29). We cloned the dIno80 motif upstream of the promoter (BS-up) and downstream of the poly-A signal (BS-dn) of the luciferase gene as described previously (29) (Fig. S3). The BS-dn clones were generated as PcG and Trx complexes are known to affect transcription when present either upstream or downstream of the target gene and dIno80 as an ETP (Enhancer of Trithorax and Polycomb) protein interacts with both the complexes (23–24, 37).

The pGL3, BS-up and BS-dn plasmids were transfected into S2 cells and luciferase activity was measured. We observed a significant up-regulation (6.88 fold) of luciferase activity in pGL3-BS-up relative to the expression from the empty vector (Fig. 6, p-value=0.00017, Student’s T-test). To examine, the specificity of this regulation, dIno80 was knocked-down with siRNA, which led to the reversal of reporter gene expression (0.71 fold, p-value= 0.0004, Student’s T-test) to the levels similar to pGL3 control (Fig. 6). The presence of the motif downstream to the reporter gene showed marginal down-regulation (0.87 fold, p-value= 0.13, Student’s T-test) of luciferase reporter expression, which was not found to be significant. However, on knock-down of dIno80, the expression from BS-dn construct showed low (1.6 fold increase, p-value= 0.000041, Student’s T-test) but significant increase in luciferase activity. The two different control siRNA duplexes were unable to affect the reporter gene expression, thus confirming dIno80 dependent regulation of the reporter. In contrast to hINO80, DBINO domain, the dIno80 DNA binding activity led to transcriptional activation. Thus, the presence of dIno80 binding motif, both upstream as well as downstream of the reporter gene, can regulate reporter gene expression, but with different outcomes.

**FIGURE 6.**
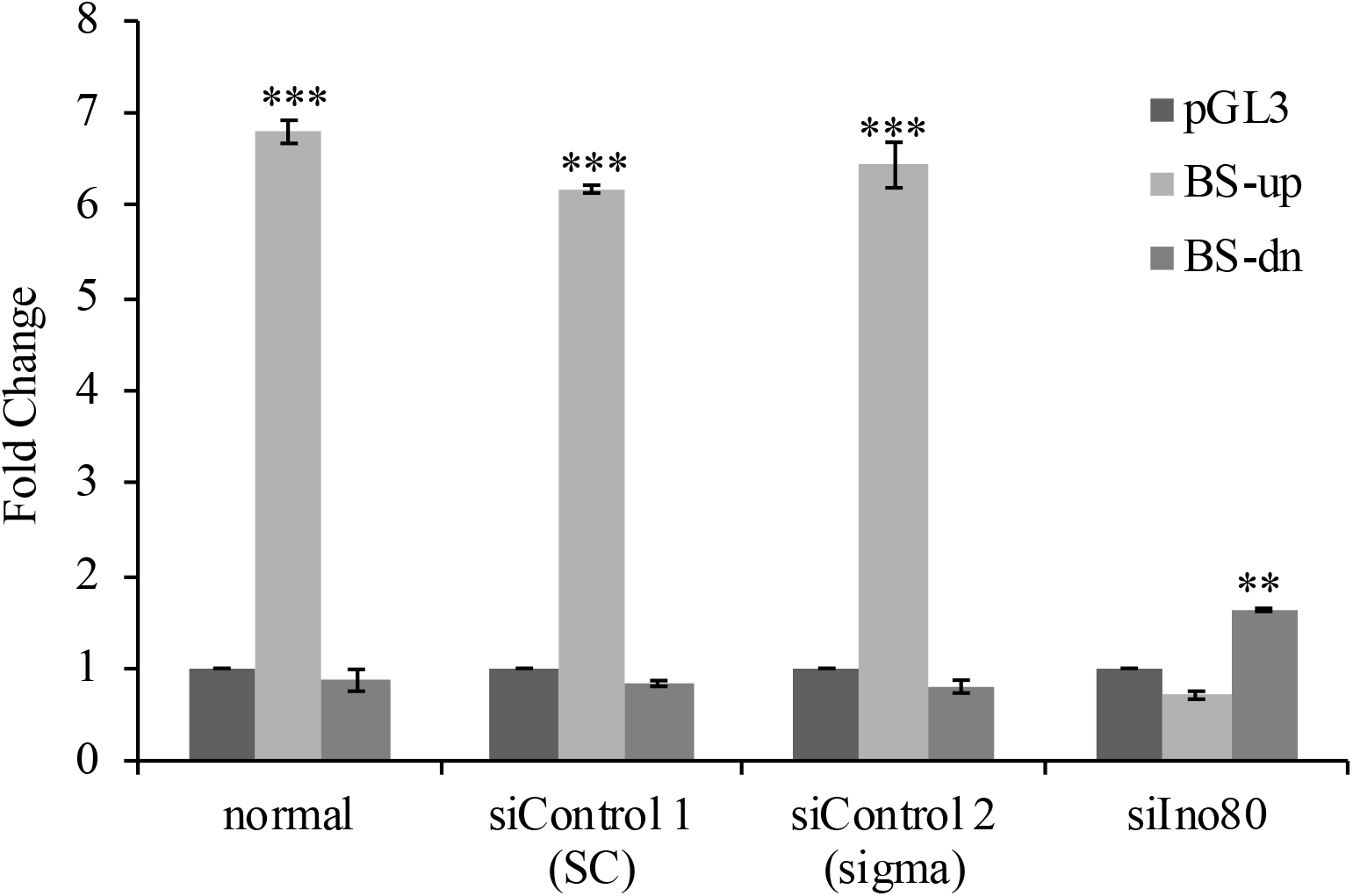
Effect of Ino80 binding motif (Hs.con) on the expression of reporter gene in S2 cells. The expression of the reporter, firefly luciferase from the constructs is indicated as fold change with reference to the expression from pGL3 vector. The effect of Ino80 binding on the expression of reporter gene from pGL3 with Hs.con under normal and on knock-down of Ino80. (** p-value <0.005, *** p-value <0.0001).

### Variable affinity of *Drosophila* INO80 for DNA

The Hs.con was derived using the DBINO domain of the human homologue of INO80 as the bait in SELEX assay (29). The DBINO domain of dIno80 and hINO80 are largely similar, but there are a few variations in the amino acid sequence. Therefore, we tested variant oligonucleotides for their interaction with dIno80, replacing one/two nucleotide(s) at a time. We evaluated the relative binding affinity of DBINO domain for the variant sequences using the FID assay. All the variants, except V6, had higher affinity than Hs.con (Table 1). The specificity of this interaction was demonstrated by the failure of the random sequence to displace EtBr in FID assay.

**Table 1.** DBINO domain interacts with Hs.con and its variants. The affinity of DBINO domain to Hs.con and the variant oligoes. The variants of human consensus INO80 binding motif (Hs.con) show higher affinity for the *Drosophila* DBINO domain as seen by fluorescence spectroscopy results. The variants are the mutants of Hs.con sequence and are same as M1, M2, M3, M4, M5, M6 and M12 (Mendiratta *et al*., 2016).

| Double stranded oligonucleotide | Sequence | Apparent $K_d$, μM |
| --- | --- | --- |
| Hs.con | GTCAGCC | 10 |
| V1 | GTCAGTT | 0.5 |
| V2 | GTCATCC | 0.6 |
| V3 | GGCAGCC | 0.8 |
| V4 | TTCAGCC | 0.5 |
| V5 | GTCCGCC | 5 |
| V6 | GACAGCC | 8 |
| V12 | GTCAGTC | 2 |

We further examined the interaction of the dIno80 in nuclear extract with the variant sequences. The variants were used as cold competitors against Cy3 labeled Hs.con oligonucleotide (CCCCGTCAGCC; Fig. 7 A). The increasing molar ratio of labeled to unlabeled probe was used (1:1, 1:2 and 1:5). All the variant sequences, except V7, displaced Hs.con at 1:5 ratio and some at lower ratios. The variant V1 (CCCCGTCAG**TT**) and V3 (CCCCG**G**CAGCC) showed higher affinity for DBINO domain of dIno80 as they almost completely displaced the Cy3-Hs.con oligo at 1:2 molar ratio. The variant V7 (CCCCGT**T**AGCC) was unable to compete with the Hs.con motif for binding unlike the other variant sequences (Fig. 7 B).This reflected the importance of certain positions within the motif for interaction. To confirm the higher binding affinity of dIno80 to the variant oligoes, we performed a competitive EMSA with Cy3 labeled V1 oligonucleotide at 1:5 molar ratio (probe: protein) and used Hs.con as the competing oligo (Fig. S4). The V1 variant, even at 1:10 ratio, was only marginally competed out by Hs.con. This further confirmed that the dIno80 protein recognized variants of the Hs.con sequence with different affinity, in concordance with the spectroscopic analysis carried out with purified DBINO domain. Thus, dIno80 has differential affinity for variants of Hs.con, which can result in dynamic interaction of the protein at different genomic positions depending on the abundance of dIno80 in the tissue. This is discussed further in the later section.

**FIGURE 7.**
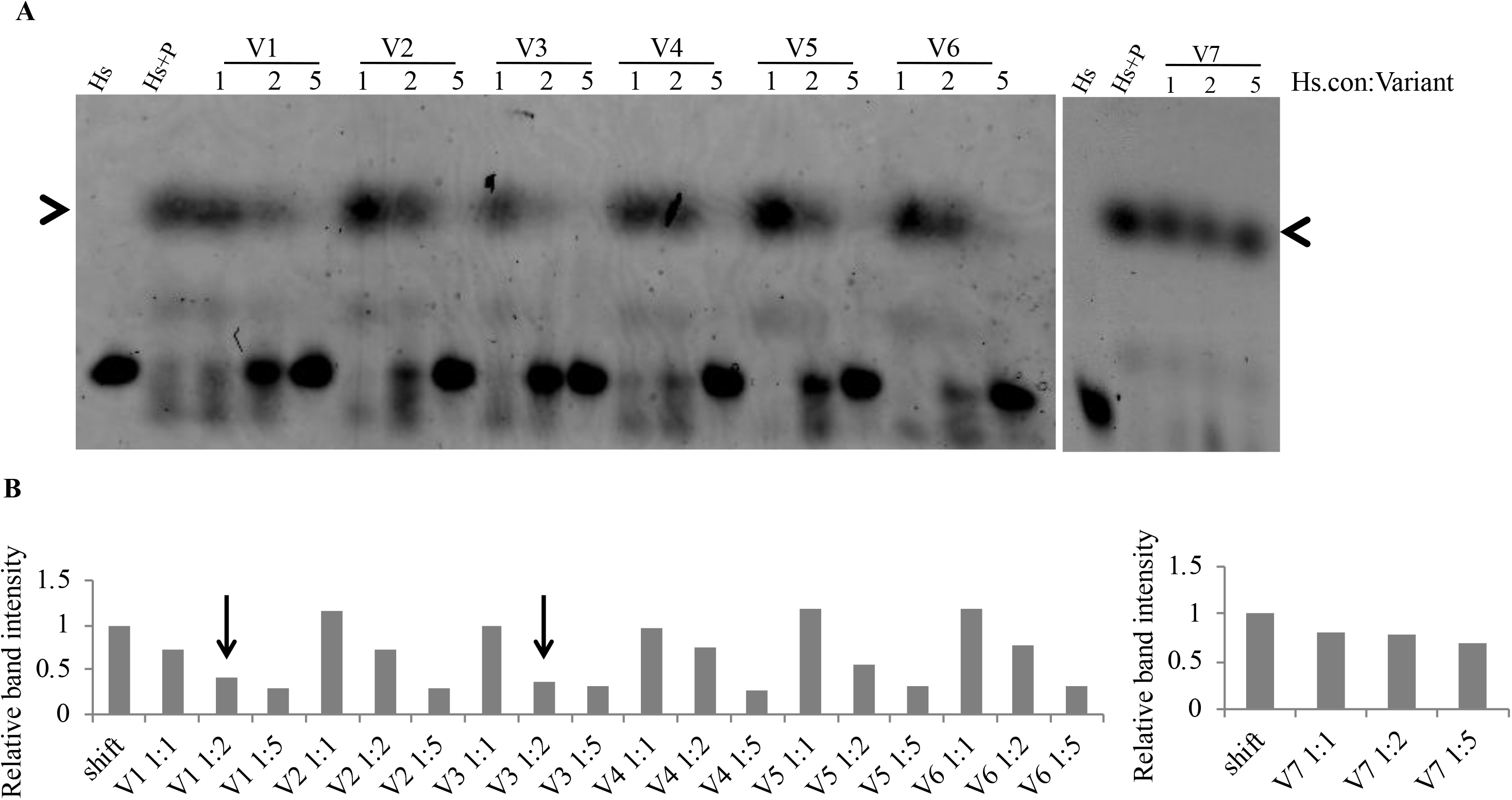
Competitive EMSA with the variant oligonucleotides (V1-V7) and Hs.con. (A) Cy3 fluorescence captured by phosphoimaging. (B) Densitometric scan of the gel. The relative intensity is calculated as the ratio of the intensity value of the retarded band (arrowhead on the gel image) in presence of the competing oligo normalized to band at the same position in lane Hs+P (1:5) without the competing oligo. Three different ratios (1:1, 1:2, 1:5) Hs.con:competing oligo was used at as indicated on top of the lanes.

In order to investigate the effect of variant sequences on the regulatory function of dIno80, these sequences were cloned in pGL3 and luciferase expression was assayed (Fig. 8, A-C). The pGL3 constructs containing the variants V1-V4, both upstream and downstream of the reporter gene were used in these assays. We observed that the up-regulation of reporter expression with variant sequences was higher than with Hs.con motif (Fig. 8, A and B). These results correlated with the results of the spectroscopy and EMSA experiments, dIno80 had higher affinity for the variants of the human consensus motif. In comparison to Hs.con, reporter expression was significantly different with the variants V1 and V4 when cloned upstream (Fig. 8 A), while all four variants led to increased expression when cloned downstream (Fig. 8 B). The V1 variant showed the highest effect on the reporter gene expression, which corresponds to the *in vitro* interaction results. The specificity of this reporter gene expression was also tested by knocking down dIno80 expression using siRNA duplexes. The knockdown of dIno80 led to lower expression of dIno80 and hence, reversal in the up-regulation of the reporter gene expression to the basal level as in the control vector (Fig. 8 C). The reason for the difference in the reporter expression in BS-dn clones of Hs.con and the variant sequences was not clear at this stage.

**FIGURE 8.**
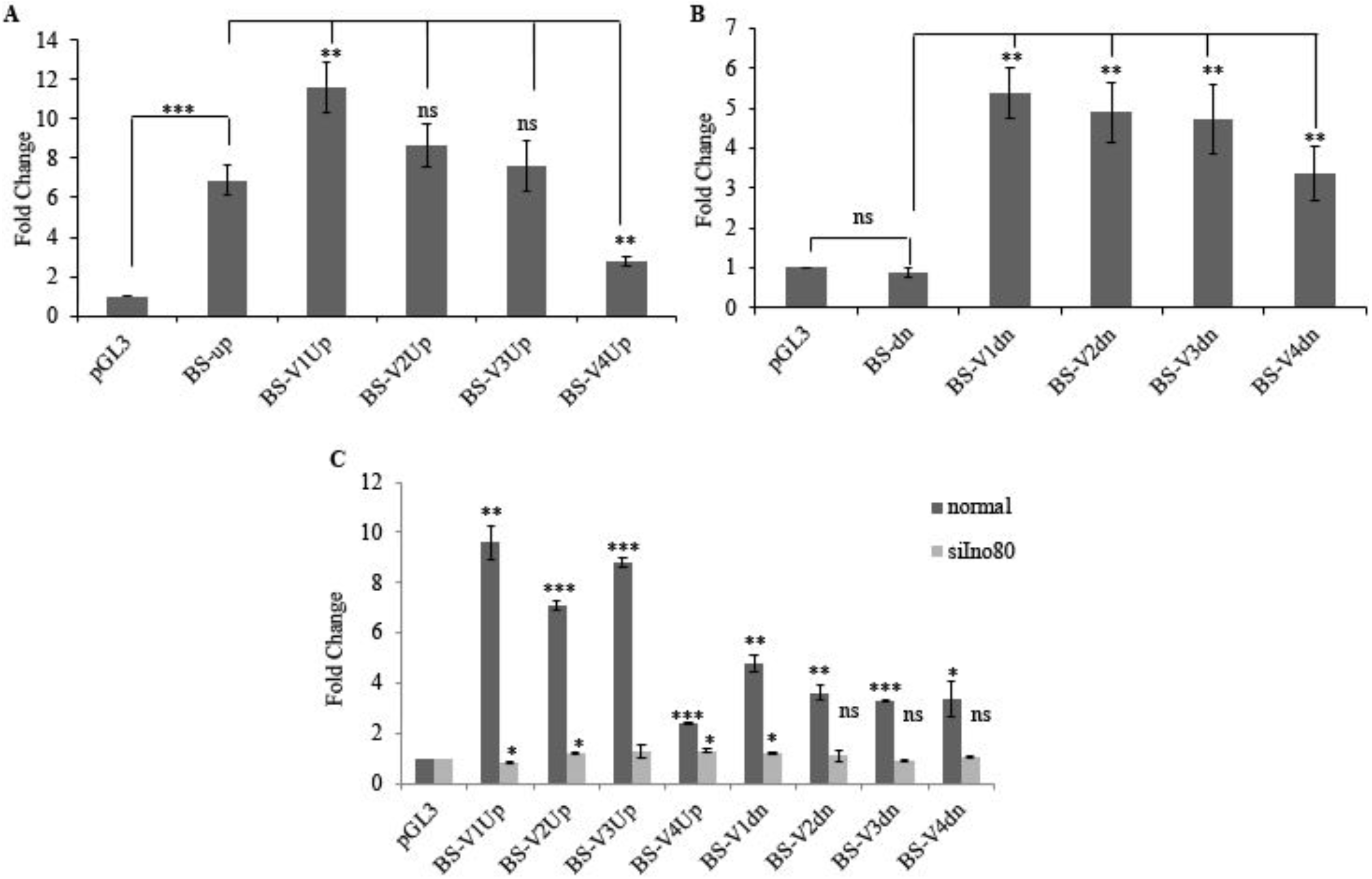
Effect of Hs.con and its variants on the expression of reporter gene in S2 cells. The expression of the reporter, firefly luciferase, from the constructs is indicated as fold change with reference to the expression from pGL3 vector. (A) and (B) The effect of the variant oligos V1-V4, on reporter expression. (C) The specificity of Ino80 binding on the expression of luciferase reporter gene in S2 cells. The activation of reporter expression is dependent on the presence of dIno80 in the cells. BS-binding site, Up-Upstream, Dn-downstream. For more details please see the text. The error bars represent the standard deviation (*p-value <0.05, ** p-value <0.005, *** p-value <0.0001, ns-not significant).

### Genome-wide analysis of dIno80 and pho binding sites

To examine the correlation of Ino80 binding and the occurrence of DNA sequence motif for Ino80, we analyzed the ChIP data for dIno80 and Pho protein available in the public domain. The dIno80 ChIP-chip data was retrieved from Moshkin *et al*., (33) and Pho data from Schuettengruber *et al*., (34). These data were analyzed to identify exclusive dIno80/Pho bound regions and also the shared binding regions. We carried out a motif search using MEME-ChIP (38) on these sequences and detected a sequence highly similar to Hs.con as the second most frequently occurring motif with an e value of 2.9e-10 and with a frequency of occurrence of 4077/7590 sites (C[TA]GCT[GC]C). We have analysed the occurrence of dIno80 binding motif in the regions identified as dIno80 binding sites in the ChIP experiments (34). Thus, the query sequences were limited to 300-500bp, as obtained in the ChIP experiments. The occurrence of Ino80 binding motif was analysed with these regions. In a second step, we analysed the occurrence of genes within ±2000bp relative to the motif recognized. This length was chosen as in a previous analysis, we observed that the difference in nucleosome organization relative to TSS (Transcription Start Site) in active and inactive genes decrease around ± 1000bp (39). We found three classes of targets: 4677 exclusively for dIno80, 1691 exclusively for Pho and 1410 genes as common targets for both Pho and dIno80 (Fig. S5). The gene ontology classification of these target genes was performed using PANTHER v12 (data not shown). When classified on the basis of biological processes, most of the target genes were enriched in categories such as cell communication, cell cycle and metabolic processes. Another GO term found enriched was cellular biogenesis. This highlighted the role of dIno80 in cellular homeostasis and development. The primary function for Pho protein in the Pho-dIno80 complex is to act as a recruiter of the repressive protein complex to the site of action on chromatin through sequence specific interaction with the DNA (19). The frequency of occurrence of exclusive dIno80 binding sites on the genome points towards the Pho independent function of the complex where dIno80 can interact as a recruiter protein to direct the complex (canonical or non-canonical) to the site of action, even if the sequence dependent recruitment of Pho is not available at these sites.

## DISCUSSION

INO80 is well-known for its functional versatility participating in multiple processes, including DNA damage repair and transcription regulation (27, 40). The chromatin remodeling proteins are rarely reported to contain a sequence specific DNA binding domain. However, CHD1, a chromatin remodeling protein, is known to have a DNA-binding domain similar to SANT-SLIDE domain of ISWI (41). Though, CHD1 does not exhibit sequence specific DNA binding activity. Here, we report the DNA binding activity of dIno80 protein. The DBINO domain showed sequence specific DNA binding activity and regulates transcription and is highly conserved within the INO80 subfamily of chromatin remodeling proteins (16–17, 29). This domain overlaps with the HSA domain which is known to interact with actin related proteins, ARPs. The X-ray crystal structure of the Arp8 and Arp4 module of yeast INO80 showed that the DBINO/HSA domain forms a helical structure (42). The HSA domain is crucial for nucleosome sliding but not for binding and ATP hydrolysis (42). The structure and interaction of the sub-complexes of the canonical INO80 complex has been analysed; it was shown that Ino80 (via the HSA domain), Arp8 (via N-terminus) and Arp4 (via C-terminus) interact with the extra-nucleosomal DNA in yeast (43). The presence of a domain named HSA (helicase-SANT associated) at the N-terminal of hIno80 has also been described (44). The SANT domain is designated after the different proteins (<u>S</u>WI3, <u>A</u>DA2, <u>N</u>-CoR and <u>T</u>FIIIB) sharing the domain (45). The SANT domain has no reported DNA binding activity and is known to interact with histone tails (46). The HSA domain overlaps with the DBINO domain, and is required for nucleosome remodeling activities through its interaction with Actin, Arp8 and Arp4 (21, 47). These reports show that HSA domain is the binding module for the ARP proteins which in turn contacts DNA to bring about chromatin remodeling (21, 48). These are most likely the canonical complexes of INO80 and much more abundant in the cell. However, non-abundant INO80 complexes lacking the ARP module (possibly the non-canonical form), may rely on DNA binding activity of DBINO domain to bring about transcriptional regulation. This is relevant in the frame work of our hypothesis wherein we propose that INO80 interacts with different complexes generating non-canonical complexes, in addition to the canonical complex.

The affinity of the DBINO domain to DNA especially with the Hs.con is low; however, we observed that compared to the cloned protein, the nuclear extracts showed higher affinity to DNA (3μM). The human and yeast Arp8 has been shown to bind to DNA with low affinity, while Arp4 showed no interaction with DNA (36). The sub-complex showed substantial DNA-binding activity in 366nM range, which was attributed to DBINO/HSA of INO80 and minor contribution of Arp8 (36). The affinity of the variant sequences that we have analysed varies from 500nM to 8000nM; V1 and V4 show affinity of 500nM, which is in similar range as estimated for the sub-complex of INO80 and Arp (36).

The interaction of dIno80 with variable motifs within the consensus sequence is particularly relevant to developmental regulation. This differential affinity suggests one of the possible ways in which the developmental genes are regulated in a spatial and temporal context. The varying protein gradient and the differential binding affinity of trans factors to the cis-elements have a key role in transcriptional regulation of target genes across the body axis (49–50). Crocker *et al*., (51) reported that clusters of low affinity binding sites for Ultrabithorax protein are needed for specifically regulating *shavenbaby* gene in *Drosophila*. De *et al*., (52) have shown that deletion of four strong PREs in the *invected-engrailed* (*inv-en*) gene complex led to no change in expression patterns of PcG regulated genes in embryo and larvae. They identified three weak PREs in *en-*fragment which can function to form a Polycomb domain to regulate gene expression (52). Stathopoulos *et al*., (53) have shown that *Dorsal* gradient formation in *Drosophila* embryo is essential to specify dorsal-ventral axis. This gradient is maintained by binding sites with variable affinity upstream of the *twist* gene. Such variable affinity of binding sites in *Drosophila* species has been shown to lead to the segmental expression patterns (54). Kwong *et al*., (55) have identified many variable binding sites for Pho in the *Drosophila* genome, which partially overlap with Polycomb binding sites. Loubiere *et al*., (56) have shown the presence of weaker Pho binding sites which are associated with transcriptional activation mediated by PRC1 complex exclusively. The binding of Pho and dIno80 independent of each other is reflected in the detection of exclusive motif for Pho and also dIno80 binding, in addition to shared sites. This suggests that these two proteins can form different complexes other than the canonical INO80.com as in the model we suggest in the context of hIno80 (29) and the non-canonical INO80 complex reported (57).

Similar variation in binding sequence is known for YY1, the human homologue of Pho (58). The genes downstream to YY1 have a variant of the known motif (CGCCATnTT); GGCGCCATnTT (Peg3) and CCGCCATnTT (*Xist*). Further, the longer motif recognized by YY1 (GCCGCCATTTTG), has higher affinity for the protein, similar to our observation of higher affinity of dIno80 to the variant sequences. In case of YY1, this is implicated in other functions of YY1 important for recruitment of PcG complexes but not in transcriptional repression (59).

The binding affinity of Hs.con with the DBINO domain is low compared to that of the variant sequences. However, Hs.con shows higher affinity with nuclear extract, which can be attributed to the difference between the shorter DBINO domain and the full length dIno80 protein present in the nuclear extracts, hence the structural stability. Further, we note that the spectroscopic method used involves the fluorescence detection of external dye which gives the apparent K_d_ (60). More accurate methods like, isothermal calorimetry can be deployed to get the binding affinity more precisely. The higher affinity of the variant sequences in competition EMSA with nuclear extract is concordant with the increased affinity measured by FID assay. The higher affinity of the variants is also reflected in their effect on reporter expression.

It is suggested that dIno80 interacts with the genome to bring about transcriptional regulation (23). INO80 complex is recruited to promoters of pluripotency genes by its interaction with transcription factors OCT4 and WDR5 (28). In cervical cancer cells, it was reported that INO80 localized at the TSS of Nanog, promoting its expression, and leading to cell proliferation (57). The localization of chromatin remodeling proteins can lead to epigenetic changes, while their own recruitment may be through recognition of modified histones. Ino80 was found to be colocalized with active histone marks H3K4me3 and H3K27ac on the TSS of pluripotency genes like OCT4, NANOG and SOX2 (28). We have reported colocalization of hIno80 with repressive histone mark, H3K27me3 in the upstream region of HOXC11 and PAX7 in HEK cells, where hIno80 binding motif (Hs.con) is present (28).

The human INO80 complex has been shown to form two distinct complexes-canonical and non-canonical complex in liver cancer cell line (61). Most of the canonical Ino80 complex peaks were found proximal to TSS of highly expressed genes colocalizing with H3K27ac histone mark. The much less abundant non-canonical Ino80 (NC-INO80) peaks were found colocalizing with repressive chromatin marks, H3K27me3 and EZH2. Though, there was no direct physical interaction with PRC2, NC-INO80 are distinctly sites of transcription repression. Another non-canonical Ino80 complex has been reported in yeast and murine embryonic cells, called as MINC complex (62). In this complex, Ino80 partners with transcription regulators MOT1 and NC2, which are known transcriptional suppressers (63). Our results with hINO80 and dIno80 evoke an additional function for Ino80 protein and imply that it can function as a recruiter of regulatory complexes. Further, the occurrence of the variant sequence motifs in the genome upstream and downstream of protein coding genes in *Drosophila* is significantly high (data not shown). The analysis of the ChIP-chip data of Moshkin *et al*., (33) has shown the localization of dIno80 at multiple sites in the genome. This suggests that Ino80 can be part of multiple complexes and depending on the interacting proteins, the outcome of interaction could either have a positive or a negative effect on transcription. However, it remains to be seen if Ino80 functions as recruiter by virtue of its DNA binding ability in these cases.

We have previously characterized dIno80 as an ETP protein (Enhancer of Trithorax and Polycomb) that can interact with PcG as well as Trx complex in *Drosophila melanogaster* (23–24, unpublished data). The effect of dIno80 binding upstream as well as down-stream of the reporter gene has a significant effect on reporter expression and this is specific to the presence of dIno80 as shown by knock-down experiments. The variable effect on reporter expression can be viewed in the context of dIno80 as an ETP protein. The protein interaction in these cases remains to be investigated to explain such contrasting effect in a transient expression model.

## CONCLUSIONS

In summary, we have identified sequence dependent interaction of dIno80 with DNA and being an essential protein for the completion of development in *Drosophila*, its interaction with DNA is an essential functional attribute of Ino80. The presence of two different functional domains, DNA dependent ATPase and sequence specific DNA binding domain, in Ino80 is a unique feature of this sub-family of chromatin remodelers (10, 23). The involvement of INO80/dIno80 in a wide variety of processes can be attributed to the different interactions it can have by virtue of multiple domains. In the context of the well-known role of morphogen gradients in axis determination and segmentation pattern in *Drosophila*, it is of interest that dIno80 interacts with variant sequences around a consensus with differential affinity. This could have an important biological consequence that can correlate the occupancy of the protein at a given site to its affinity, which in turn would be sensitive to the abundance of the protein within the given cell.

### AUTHOR CONTRIBUTIONS

VB conceptualized and conceived the project and supervised the study; SJ executed the project; SM designed the spectroscopic studies and interpreted the spectra; JM contributed to experiments with S2 cells and mass spectrometry; AN helped with *in silico* analysis; VB and SJ interpreted the results and wrote the manuscript.

## ACKNOWLEDGMENTS

We thank Prof. Yogendra Singh, Dr. Gunjan Arora and Dr. Debasis Dash of CSIR-Institute of Genomics and Integrative Biology for cloning vector and Bioinformatic tools, Dr. Suman Thakur and Dr. Rakesh Mishra of CSIR-Centre for Cellular and Molecular Biology for Mass spectrometry and S2 cells respectively. This work was supported by a grant from the Department of Biotechnology, Govt India (BT/PR4442/PID/6/626/2011) and grants from the Council for Scientific and Industrial Research (CSIR), Government of India (EpiHeD:BSC0118/2012-17) and No.60(0102)/12/EMR-II). We acknowledge DBT Bioinformatics facility at ACBR and the UGC SAP-II support. SJ received a fellowship from CSIR, Government of India, JM received Research Associate fellowship from SERB EMR/2016/000653/dt 15/03/2017) and AN from D. S. Kothari Fellowship from UGC.

## Abbreviations

dIno80: *Drosophila* Ino80
hIno80: human Ino80
Ino80: Inositol requiring mutant 80
ETP: Enhancer of Trithorax and Polycomb
FID: fluorescence intercalator displacement
DBINO: DNA binding domain of Ino80
EMSA: electrophoretic mobility shift assay.

