## Supplementary material for "The regulatory function of dIno80 correlates with its DNA binding activity": Supplentary Figures

**Supplemental Figure S1**

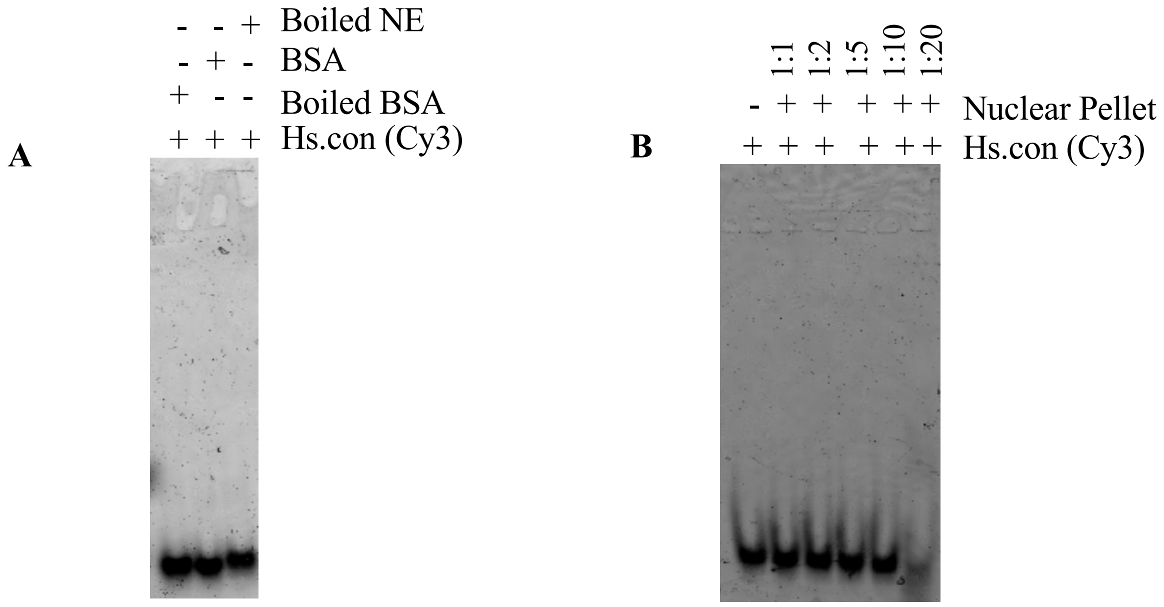

**Supplemental Figure S2**

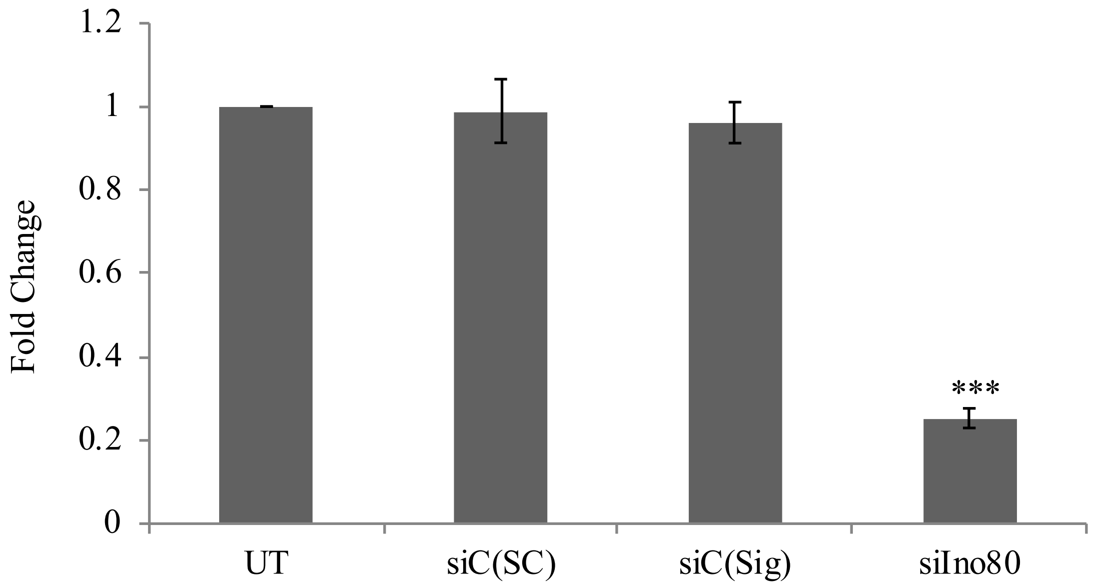

**Supplemental Figure S3**

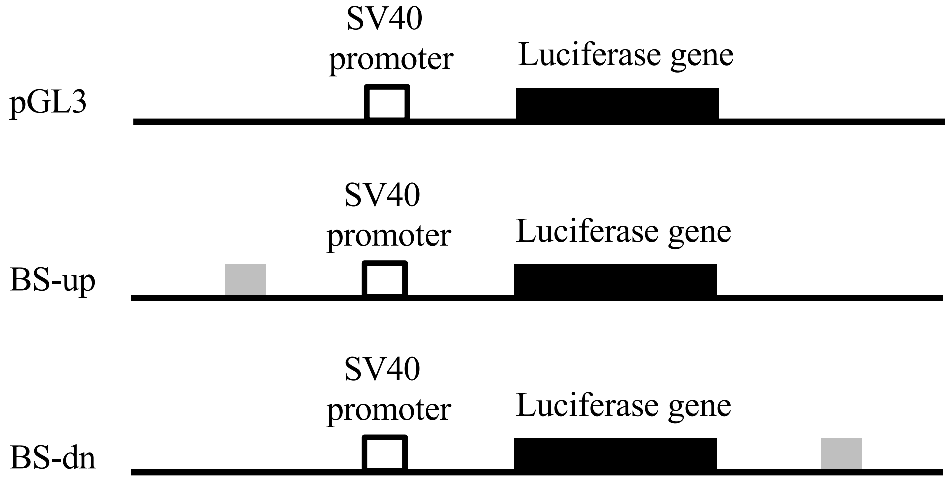

**Supplemental Figure S4**

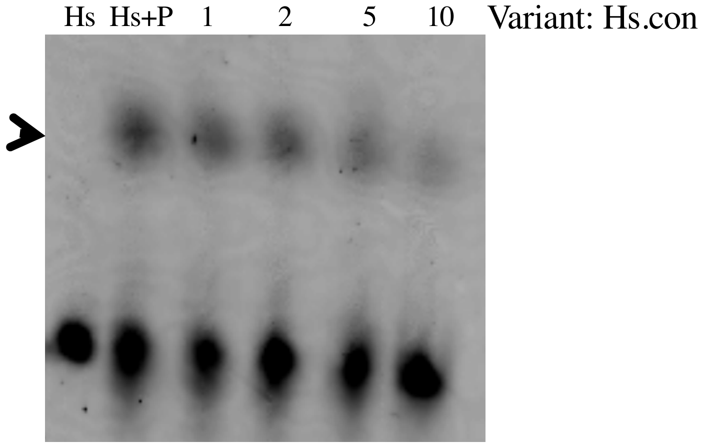

### Supplemental Figure S5

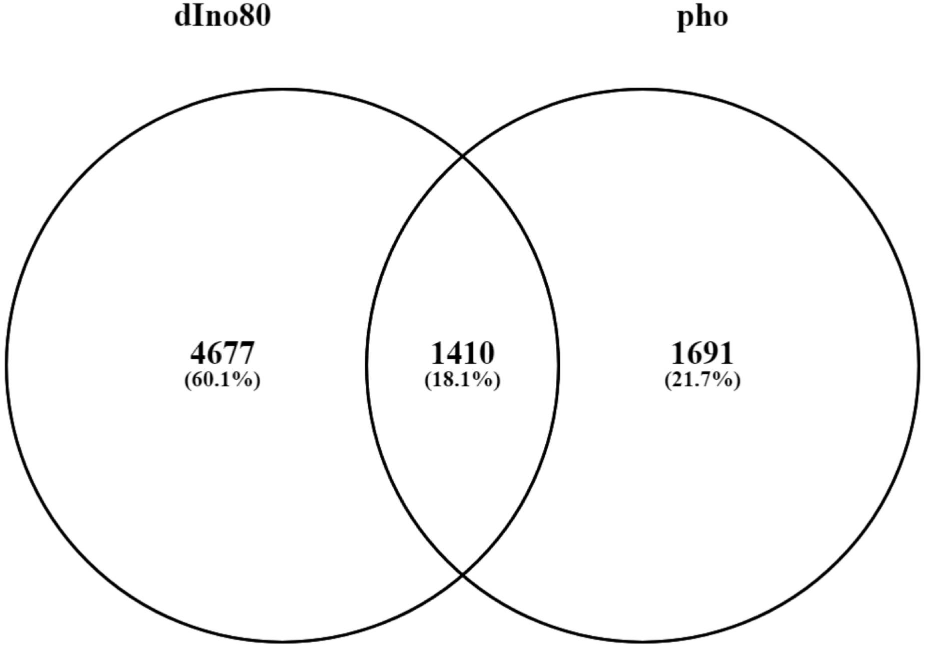
