## Supplementary Figure legends for "The regulatory function of dIno80 correlates with its DNA binding activity"

**Supplemental Figure S1**. (A) Absence of interaction of Hs.con with boiled nuclear extract from wild type larvae and bovine serum albumin (BSA) at 1:5 probe:protein ratio. (B) Absence of interaction with protein in nuclear pellet indicating specificity of interaction. The DNA amount used was 1 picomole of dsDNA at varying probe: protein molar ratio. NE-Nuclear Extract.

**Supplemental Figure S2**. The knockdown efficiency of siIno80 using the quantitative PCR. The expression of *dIno80* was normalized to that of rpl*32* expression (p<0.0001). Nearly 80% decrease is observed.

**Supplemental Figure S3**. A line diagram of the constructs used in the luciferase reporter assays. The pGL3 construct is the pGL3 promoter vector without the INO80 binding site (Hs.con, grey square), BS-up construct is where the Hs.con is cloned upstream of the promoter of luciferase reporter gene and BS-dn is where binding motif is cloned down-stream of the poly(A) signal. The constructs have been described in Mendiratta *et al*., 2016.

**Supplemental Figure S4.** Competitive EMSA with labeled variant oligonucleotide. The variant oligonuclotide V1 was Cy3 labeled and used at 1:5 (probe: protein) molar ratio. The unlabeled human consensus oligo was used as competitor at increasing ratio relative to V1 as indicated on the top of the lanes.

**Supplemental Figure S5**. The dIno80 and Pho binding site analysis. The binding peaks for dIno80 and Pho were retrieved from Moshkin *et al*., 2012 and Schuettengruber *et al*., 2009 respectively. The overlapping regions were identified and genes mapping ±2000bp within the binding sites were retrieved. These gene lists' were compared using Venny to identify dIno80 exclusive, Pho exclusive and overlapping targets.
