## Supplementary Table for "The regulatory function of dIno80 correlates with its DNA binding activity"

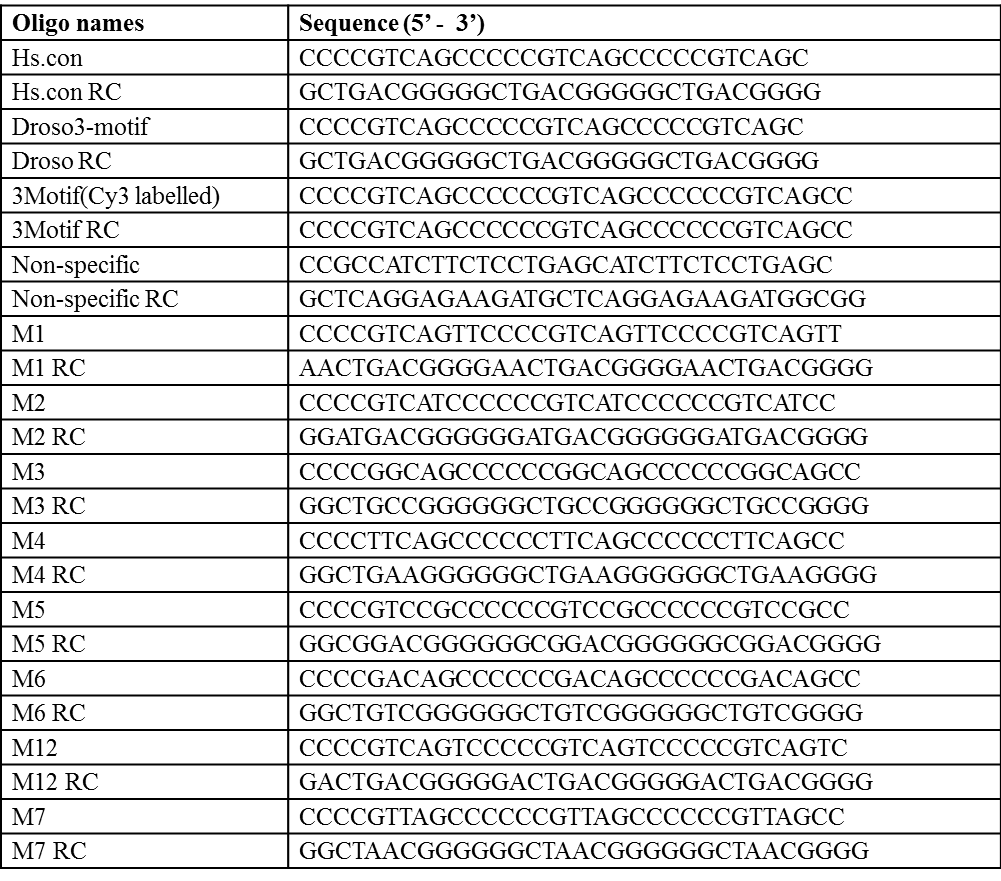

**Supplemental Table S1.** The sequences of oligonucleotides used in FID and EMSA experiments.

**SUPPLEMENTAL TABLE S1**
